# Phosphorylation of a Cleaved Tau Proteoform at a Single Residue Inhibits Binding to the E3 Ubiquitin Ligase, CHIP

**DOI:** 10.1101/2023.08.16.553575

**Authors:** Cory M. Nadel, Kristin Wucherer, Abby Oehler, Aye C. Thwin, Koli Basu, Matthew D. Callahan, Daniel R. Southworth, Daniel A. Mordes, Charles S. Craik, Jason E. Gestwicki

## Abstract

Microtubule-associated protein tau (MAPT/tau) accumulates in a family of neurodegenerative diseases, including Alzheimer’s disease (AD). In disease, tau is aberrantly modified by post-translational modifications (PTMs), including hyper-phosphorylation. However, it is often unclear which of these PTMs contribute to tau’s accumulation or what mechanisms might be involved. To explore these questions, we focused on a cleaved proteoform of tau (tauC3), which selectively accumulates in AD and was recently shown to be degraded by its direct binding to the E3 ubiquitin ligase, CHIP. Here, we find that phosphorylation of tauC3 at a single residue, pS416, is sufficient to block its interaction with CHIP. A co-crystal structure of CHIP bound to the C-terminus of tauC3 revealed the mechanism of this clash and allowed design of a mutation (CHIP^D134A^) that partially restores binding and turnover of pS416 tauC3. We find that pS416 is produced by the known AD-associated kinase, MARK2/Par-1b, providing a potential link to disease. In further support of this idea, an antibody against pS416 co-localizes with tauC3 in degenerative neurons within the hippocampus of AD patients. Together, these studies suggest a discrete molecular mechanism for how phosphorylation at a specific site contributes to accumulation of an important tau proteoform.

## INTRODUCTION

The deposition of microtubule-associated protein tau (*MAPT*/tau) into insoluble, neurofibrillary tangles (NFTs) is a major pathological hallmark of fatal and incurable neurodegenerative diseases – including Alzheimer’s disease (AD), frontotemporal dementia (FTD), progressive supranuclear palsy (PSP), corticobasal degeneration (CBD), and others^1-3^. There is interest in understanding the pathways that govern tau proteostasis, in both health and disease^4^, because such knowledge may reveal mechanisms for reducing the accumulation of NFTs.

Many studies have suggested that CHIP (C-terminus of Hsc70 interacting protein; *STUB1*) is one of the major E3 ubiquitin ligases responsible for degradation of tau^5^. For example, the CHIP^-/-^ mouse accumulates tau^6^, CHIP overexpression reduces tau aggregation^7^, and a recent unbiased screen identified the CHIP pathway as a key regulator of tau pathogenicity^8^. In the canonical mechanism, CHIP is recruited to tau via the molecular chaperone, heat shock protein 70 (Hsp70)^9^. In the first step, Hsp70 binds to at least two major sites in tau’s microtubule-binding repeats (MTBRs) ^10-14^. Then, Hsp70 uses a C-terminal sequence, termed the EEVD motif, to bind CHIP’s tetratricopeptide repeat (TPR) domain^15-19^. In this way, certain isoforms of Hsp70^20^ act as adapters, recruiting CHIP and promoting tau’s ubiquitination and turnover^12,21^. However, the CHIP-Hsp70 complex does not seem to act on all tau proteoforms in the same way. Tau is found in a large number of proteoforms, including those resulting from alternative splicing and post-translational modifications (PTMs), such as phosphorylation^22^. These different tau proteoforms seem to have distinct ways of interacting with CHIP, which likely impacts their relative rate(s) of turnover. For example, the Hsp70-CHIP complex preferentially recognize pathologically phosphorylated tau^23,24^. Moreover, some tau proteoforms even bind CHIP without the need for chaperone. For example, while CHIP has relatively poor affinity for unmodified tau^25^, it directly interacts with tau that has been phosphorylated at sites in the N-terminal domain (NTD) and proline-rich region (PRR) by glycogen synthase kinase 3β (GSK-3β)^26^ Together, these studies show that CHIP interacts with different tau proteoforms using chaperone-dependent and chaperone-independent mechanisms and that these contacts are broadly sensitive to phosphorylation.

Despite these substantial advances, the molecular mechanisms that govern CHIP’s recognition of tau proteoforms are not clear. We saw an opportunity to address this question by studying tauC3, a tau proteoform that is the epitope for the widely used C3 antibody. TauC3 is the product of caspase cleavage at aspartate 421 (D421) in tau and multiple lines of evidence suggest that this truncation plays an important role in neurodegenerative disease^27^. For example, immunoreactivity with the C3 antibody strongly correlates with progression of AD^28^ and other tauopathies^29^. In addition, the cleavage event to produce tauC3 seems to precede NFT deposition^30,31^, treatment with the C3 antibody partially blocks tau seeding^32^, the tauC3 protein is more aggregation-prone than full-length tau and expression of this proteoform inhibits microtubule dynamics and slows axonal transport in neurons^33-35^. Like other tau proteoforms, tauC3 is known to be a substrate of CHIP. For example, the CHIP ^-/-^ mouse preferentially accumulates tauC3^6^ and this isoform is very rapidly ubiquitinated by CHIP *in vitro*^36^. We reasoned that the tauC3 system might be a particularly good model for studying the effects of phosphorylation on CHIP binding because the interaction site is comparatively well defined. Specifically, tauC3 has an EEVD-like motif at its C-terminus, which, similar to Hsp70, binds with high affinity to CHIP’s TPR domain^36^.

Here, we explore how tauC3 phosphorylation impacts its binding to CHIP. This question is important because, despite being an excellent substrate for CHIP, tauC3 accumulates in the brains of AD patients^28^, suggesting that unknown factors allow it to evade CHIP-mediated quality control. To explain this apparent dichotomy, we crafted a hypothesis based on data from our group and others, in which phosphorylation of a specific residue in Hsp70’s EEVD motif was found to block its binding to CHIP^37,38^. We noticed that tauC3’s EEVD-like motif has a serine at the equivalent position, S416 (numbered according to the 2N4R-tau splice isoform); thus, we hypothesized that its phosphorylation might restrict CHIP binding. Here, we apply a multidisciplinary approach to show that, indeed, pS416 is sufficient to inhibit the CHIP-tauC3 interaction *in vitro* and in cells. To understand the molecular mechanism, we used crystallography to show that pS416 creates a steric and electronic clash in CHIP’s TPR domain and we leveraged this information to create a CHIP point mutant (D134A) that partially regains the ability to bind and ubiquitinate tauC3 pS416. This mechanism might be important in disease, because we find that pS416 co-accumulates with tauC3 in dysmorphic neurons within the hippocampus of AD patient brains and that the AD-associated^39-41^ kinase, microtubule affinity regulating kinase 2 (MARK2/Par-1b), generates pS416. Together, these studies provide insight into how a hierarchical series of tau PTMs – first proteolysis to create tauC3, then phosphorylation by MARK2 to block CHIP binding and finally CHIP-mediated transfer of poly-ubiquitin -balances tau proteostasis.

## RESULTS

### Phosphorylation of tauC3 inhibits interaction with CHIP

Previous work has shown that phosphorylation of Hsp70’s C-terminus blocks binding to CHIP^37,38^. Thus, we hypothesized that a similar mechanism might govern the binding of tauC3’s C-terminal degron to CHIP. To test this idea, we first used a live cell NanoBiT split-luciferase assay^42^ to measure CHIP-tauC3 interactions in HEK293 cells (Fig 1A). As a control, we first confirmed^36^ that CHIP binds approximately 5-fold better to tauC3 over full length (FL) tau under basal conditions (Fig 1B). Throughout the manuscript, we use FL to refer to the 0N4R splice isoform of tau. Next, we treated with the protein phosphatase inhibitor, okadaic acid (30 nM), and found that it enhanced CHIP’s interaction with FL tau, consistent with the literature^23,24^, while it significantly weakened the interaction with tauC3 (Fig 1B). Thus, it seems that phosphorylation has dramatically different effects on FL tau and tauC3, highlighting the proteoform selectivity in CHIP binding. To support these cell-based observations with biochemical studies, we then purified recombinant, natively phosphorylated tauC3 (p.tauC3) protein from *Sf9* insect cells (Supplemental Fig 1A). Consistent with previous studies^23,43^, p.tauC3 produced in this way is heavily phosphorylated, as measured by immunoblots for specific tau phospho-epitopes, including pS202/pT205 (AT8), pS396 (PHF13), and pS416 (Supplemental Fig 1B). Using ELISA (Fig 1C, 1D), we confirmed that p.tauC3 binds ∼4-fold weaker to CHIP (K_d_ = 0.70 ± 0.12 μM) than unmodified tauC3 (produced from *E. coli*; K_d_ = 0.19 ± 0.02 μM). We then compared these different tauC3 proteins as substrates for CHIP in ubiquitination reactions *in vitro*. In these experiments, CHIP rapidly converted tauC3 to high-molecular weight (HMW) polyubiquitinated species; however, this process was attenuated for p.tauC3 (Fig 1E). We subsequently identified the ubiquitination sites present on the CHIP-treated tauC3 proteoforms by mass spectrometry (Supplemental Fig 1C). Interestingly, we observed an increase in the number of ubiquitination sites on p.tauC3 compared to tauC3, suggesting that CHIP might be finding weak, secondary sites when phosphorylation masks the primary site. To confirm that phosphorylation – and not a different PTM – was responsible for the observed weakening of the CHIP-tauC3 interaction, we treated tauC3 or p.tauC3 with lambda phosphatase, and then compared the binding of these proteins to CHIP using ELISAs. De-phosphorylation of p.tauC3 was confirmed via immunoblot (Supplemental Fig 1D). Under these conditions, we found that removing the phosphates was sufficient to completely rescue binding of p.tauC3 to CHIP (Supplemental Fig 1E). Together, these results indicate that phosphorylation of tauC3 weakens binding to CHIP and hinders the ability of CHIP to ubiquitinate this substrate.

**Fig 1.**
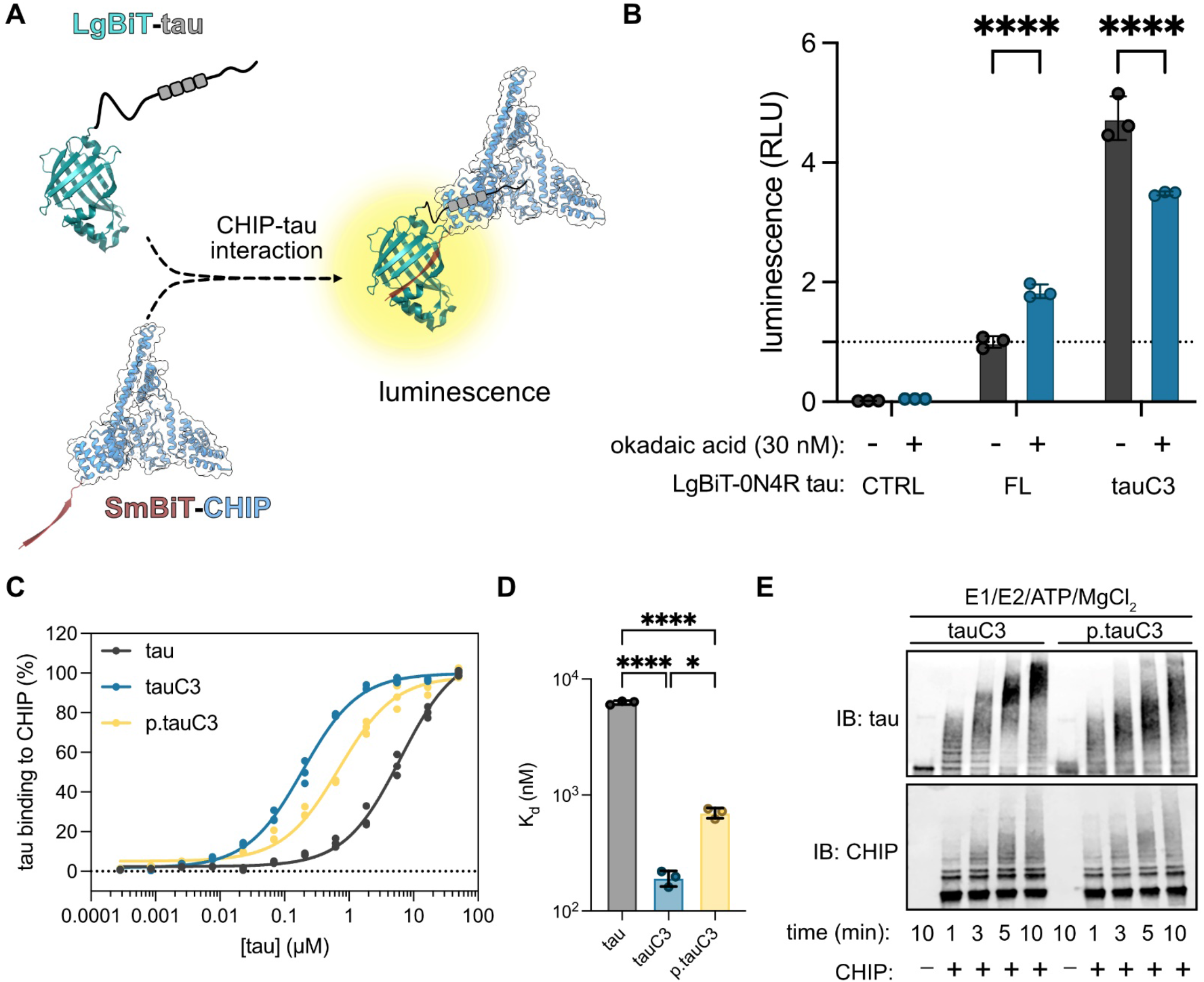
Phosphorylation of tauC3 inhibits interaction with CHIP. **(A)** Cartoon schematic of the live-cell NanoBiT assay for measuring CHIP-tau PPIs (NanoBiT PDB = 7SNX; CHIP PDB = 2C2L). **(B)** Results of NanoBiT assays for CHIP interactions with tau proteoforms following treatment with okadaic acid (30 nM, 18 hours) or DMSO control. Luminescence normalized to CHIP binding to FL 0N4R tau treated with vehicle (dashed line). Statistical significance was determined by two-way ANOVA with Bonferroni’s post-hoc analysis (****p<0.0001, n = 3). **(C)** Tau proteoforms binding to immobilized CHIP measured by ELISA. Assay was performed in triplicate and normalized to maximum absorbance @ 450 nM. **(D)** Dissociation constants derived from (C). Statistical significance was determined by one-way ANOVA with Tukey’s post-hoc analysis (*p<0.05, ****p<0.0001, n = 3). **(E)** *In vitro* ubiquitination of tau proteoforms by CHIP. Samples were collected at the denoted timepoints, quenched in SDS-PAGE loading buffer, and analyzed by western blot.

### Phosphorylation of tauC3 Ser416 is sufficient to inhibit interaction with CHIP

We next sought to identify the specific phosphorylation site(s) in p.tauC3 that might weaken binding to CHIP-tauC3. Using mass spectrometry, we identified 20 phosphorylation sites on the insect-cell derived p.tauC3 (Fig 2A). These sites were broadly located in each of the major domains of tau. Specifically, 1/20 of the sites are located in the N-terminal domain (NTD), 12/20 in the proline-rich domain (PRD), 3/20 in the microtubule-binding repeats (MTBRs), and 4/20 in the C-terminal domain (CTD). As a first step at isolating which of these sites might be important, we expressed truncated tau constructs in HEK293 cells and looked for changes in the CHIP interaction after okadaic acid treatment, as measured by NanoBiT assays (Fig 2B). Okadaic acid treatment caused nearly all the constructs to bind tighter to CHIP (Fig 2C), consistent with reports of CHIP binding to phosphorylated tau at multiple sites^26^. However, the one exception was the tauC3 construct, where, consistent with our earlier data, okadaic acid treatment inhibited binding. This result re-enforced the idea that tau proteoforms interact with CHIP through distinct mechanisms and suggested that the key phosphorylation site(s) for tauC3 are located near the C-terminus. Our mass spectrometry data showed only four phosphosites present in this region – S396, S400, S404, and S416 (Fig 2A). Due to previous work on analogous Hsp70 phosphosites^37^, we hypothesized that S416 was likely to play an important role. To test this idea, we synthesized 10-mer acetylated peptides corresponding to the C-terminus of tau (SSTGSIDMVD) and then replaced each of the possible Ser/Thr residues, including S416, with a glutamic acid as a phosphorylation mimetic. When these peptides were tested for binding to CHIP by differential scanning fluorimetry (DSF) and fluorescence polarization (FP), we found that the pS416 phosphomimetic (SSTGEIDMVD) was sufficient to block binding (Fig 2D; Supplemental Fig 2A). Specifically, by DSF, we observed an ∼6 °C stabilization of CHIP in the presence of WT tauC3 peptide compared to DMSO control (CHIP Tm_app_ DMSO = 42.15 ± 0.22 °C; SSTGSIDMVD = 48.81 ± 0.11 °C) and this binding was significantly weakened by the introduction of the phosphomimetic mutation (CHIP Tm_app_ SSTGEIDMVD = 45.49 ± 0.11 °C) (Fig 2D). In FP assays, the phosphomimetic tauC3 peptide was also a significantly weaker competitor than WT tauC3 (K_i_ WT tauC3 = 0.15 ± 0.02 μM; tauC3 S416E = 2.41 ± 0.33 μM) (Fig 2F). Critically, none of the other phosphomimetic peptides were different from WT control in either the DSF or FP platforms, indicating that only phosphorylation of the Ser416 position is important (Supplemental Fig 2B, 2C). Moreover, we introduced a *bona fide* phosphoserine residue at the S416 position and likewise observed much weaker competition by this peptide (K_i_ pS416 tauC3 = 1.01 ± 0.99 μM). To test whether this single site was important in the context of the tauC3 protein (and not just the 10mer peptide), we purified tauC3 with a single phosphomimetic mutation (S416E) and tested its binding to CHIP using ELISA (Fig 2G). Consistent with the results from the peptide experiments, tauC3 S416E bound significantly weaker to CHIP compared to WT tauC3 (K_d_ WT= 0.09 ± 0.01 μM; S416E = 0.78 ± 0.08 μM). Indeed, the effect of S416E on CHIP binding was nearly the same as the heavily phosphorylated p.tauC3 protein (see Fig 1), suggesting that this single PTM is necessary and sufficient to inhibit the CHIP-tauC3 interaction.

**Fig 2.**
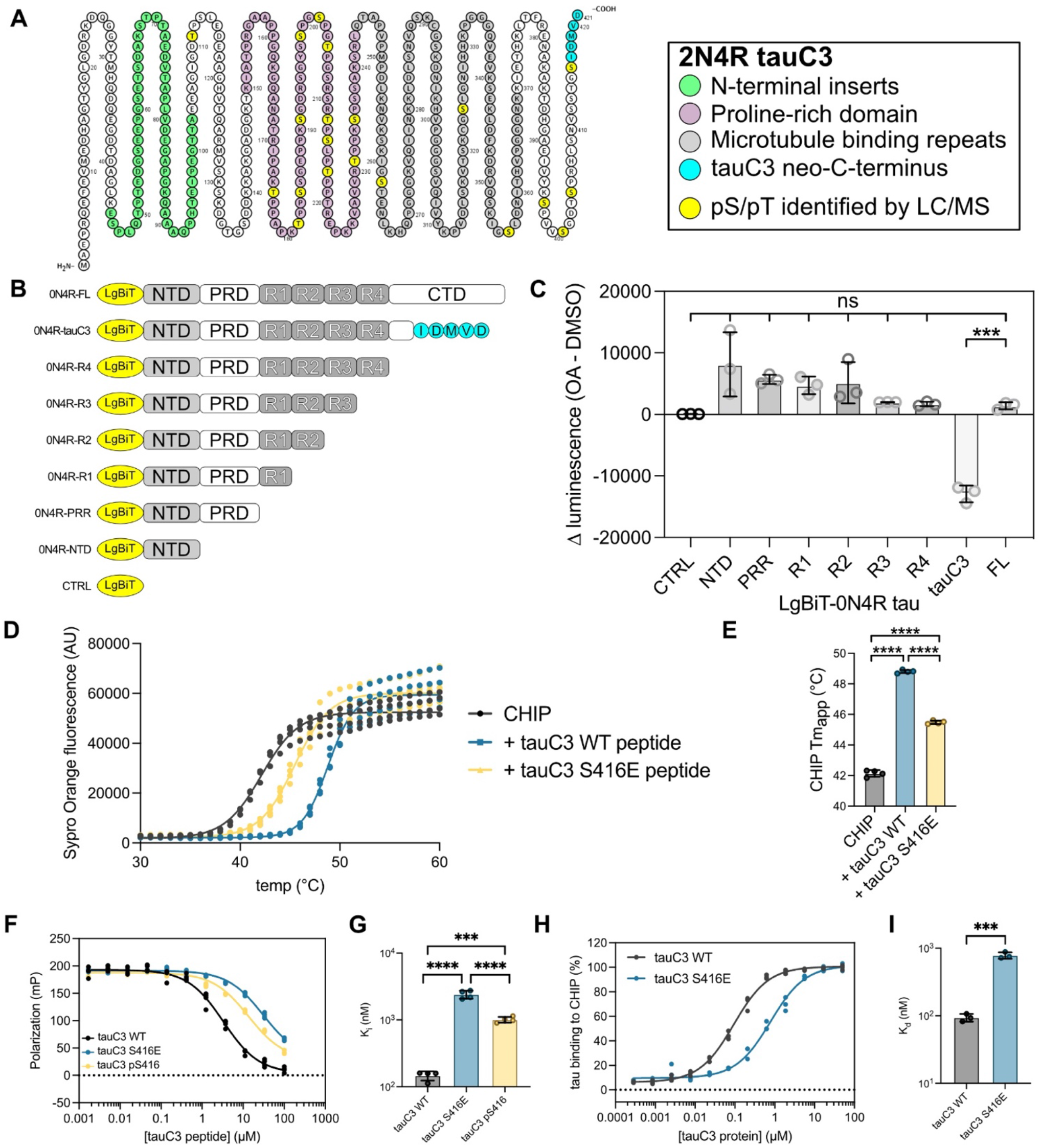
Phosphorylation of tauC3 Ser416 is sufficient to inhibit interaction with CHIP. **(A)** Locations of serine and threonine phosphorylation sites identified on p.tauC3 by LC/MS-MS. Residue numbering is derived from 2N4R tau, while identified pS/pT sites are shown in yellow. **(B)** Cartoon depicting constructs used for mapping of inhibitory phosphorylation sites by NanoBiT live cell assays. **(C)** NanoBiT live cell assay for CHIP interactions with truncated tau constructs following treatment with okadaic acid (30 nM, 18 hours) or DMSO control. Data is shown as the difference between okadaic acid treated samples and vehicle control. Statistical significance was determined by one-way ANOVA with Tukey’s post-hoc analysis (ns = not significant, ***p<0.001, n = 3). **(D)** DSF curves for CHIP incubated with DMSO control or tauC3 peptides. Assay was performed in quadruplicate and melt curves were fit with a Boltzmann sigmoid. **(E)** Apparent melting temperatures (Tm_app_) of CHIP in the absence or presence of 10-mer tau peptides as derived from (D). Statistical significance was determined by one-way ANOVA with Tukey’s post-hoc analysis (****p<0.0001, n = 4). **(F)** Competition FP experiment showing displacement of fluorescent tracer from the CHIP TPR domain by various 10-mer tau peptides. Samples were performed in quadruplicate. **(G)** Inhibition constants for various tau peptides derived from (F). Statistical significance was determined by one-way ANOVA with Tukey’s post-hoc analysis (***p<0.001, ****p<0.0001, n = 4). **(H)** Tau proteoforms binding to immobilized CHIP, as measured by ELISA. Assay was performed in triplicate and normalized to maximum absorbance @ 450 nM. **(I)** Dissociation constants derived from (H). Statistical significance was determined by unpaired student’s t-test (***p<0.001, n = 3).

### TauC3 Ser416 phosphorylation regulates CHIP-dependent tau homeostasis

We hypothesized that S416 phosphorylation might also slow CHIP-mediated ubiquitination of tauC3. To test this idea, we performed ubiquitination reactions *in vitro*. As previously observed, CHIP robustly ubiquitinated tauC3 to a greater extent than FL tau (Fig 3A,B). However, this activity was significantly attenuated by the S416E phosphomimetic mutation (Fig 3A,B). Next, we hypothesized that reduced ubiquitination might partially protect tauC3 from degradation in cells. To test this idea, we generated HEK293 Flp-In lines that express green fluorescent protein (GFP)-tagged tau variants from a doxycycline-inducible promoter (Fig 3C). Importantly, this platform involves integration of the transgene at a single shared genomic locus, reducing expression variability introduced by transfection. Using microscopy, we first confirmed that FL tau, tauC3 and tauC3 S416E all localized properly to the microtubules (Fig 3D). Next, we tested their binding to endogenous CHIP by co-immunoprecipitation. Consistent with the *in vitro* studies, we found that tauC3, but not FL tau, strongly co-immunoprecipitated endogenous CHIP from cells (Fig 3E). Moreover, this binding was partially inhibited by the S416E phosphomimetic mutation (relative amount = 0.55). In these cells, we noticed that the levels of tauC3 protein were routinely lower relative to that of FL tau or tauC3 S416E (Fig 3F,G). The reduced levels of tauC3 were not due to transcriptional effects, because the levels of each mRNA were similar, with even a modest increase in tauC3 S416E message (Fig 3H). Rather, this observation is consistent with tauC3 protein being degraded via its CHIP interaction. Together, these results indicate that the pS416 modification protects tauC3 from CHIP-mediated ubiquitination and turnover.

**Fig 3.**
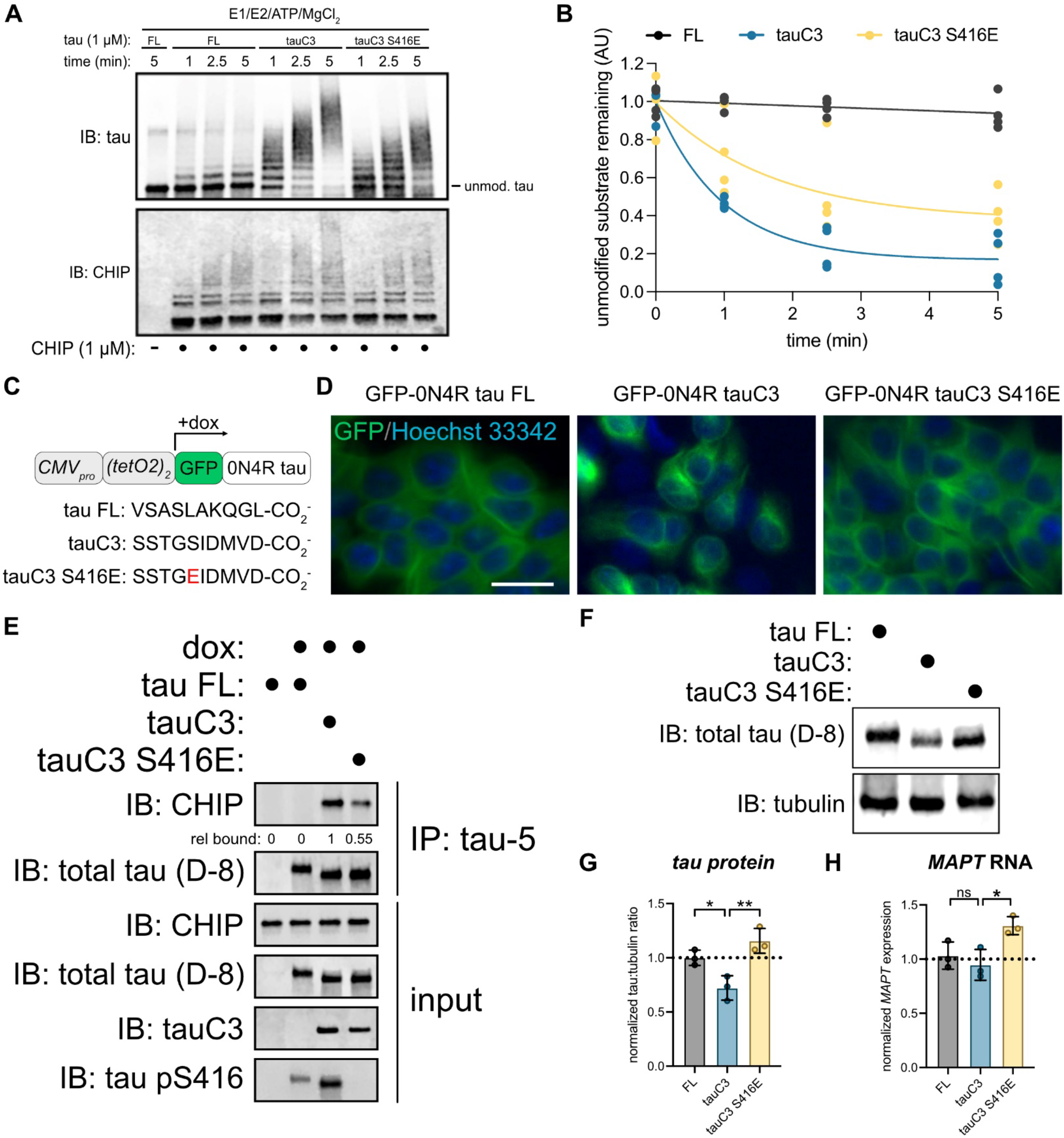
TauC3 Ser416 phosphorylation regulates CHIP-dependent tau homeostasis. **(A)** *In vitro* ubiquitination of tau proteoforms by CHIP. Samples were collected at the denoted timepoints, quenched in SDS-PAGE loading buffer, and analyzed by western blot. **(B)** Quantification of unmodified tau remaining following *in vitro* ubiquitination of varying tau proteoforms by CHIP. Unmodified tau remaining was analyzed by densitometry, normalized to 0-minute time point, and curves were fit using one-phase exponential decay (n = 3). **(C)** Cartoon schematic depicting promoter architecture and varying C-terminal sequences for HEK293 FlpIn T-Rex cells expressing doxycycline inducible GFP-tau proteoforms. **(D)** Representative fluorescence micrographs for GFP-tau cells. Images show tau species associated with microtubules (green, false color), while nuclei are stained with Hoechst 33342 (blue, false color). Scale bar = 30 μM. **(E)** Co-immunoprecipitation assay following IP of varying tau species from cells. Whole cell lysate (input) was used for loading controls, and co-immunoprecipitated CHIP was analyzed by western blot. Relative CHIP bound was determined by densitometry and normalized to tauC3. **(F)** Representative western blot showing differing abundance of various tau proteoforms in HEK293 FlpIn T-Rex cells. **(G)** Quantification of tau protein abundance taken from three independent experiments. Tau:tubulin ratio was determined by densitometry and normalized to full-length tau. Statistical significance was determined by one-way ANOVA with Tukey’s post-hoc analysis (*p<0.05, **p<0.01, n = 3). **(H)** Quantification of *MAPT* mRNA from three independent experiments. *MAPT* mRNA was normalized to *GAPDH* and shown relative to full-length tau. Statistical significance was determined by one-way ANOVA with Tukey’s post-hoc analysis (ns = not significant, *p<0.05, n = 3).

### Structural basis for CHIP binding to tauC3 and inhibition by phosphorylation

To further understand the molecular mechanism by which tauC3 S416 phosphorylation inhibits the interaction with CHIP, we solved the X-ray co-crystal structure of CHIP’s TPR domain bound to an acetylated 10-mer peptide (Ac-SSTGSIDMVD-OH) corresponding to the tauC3 C-terminus at 1.8 Å (PDB 8FYU; Fig 4A). Overall, we found that the orientation of the tauC3 peptide was similar to the “U-shaped” arrangement previously observed with other CHIP-binding peptides, with only a minor ∼1.25 Å shift of the backbone register^36,44^ (Supplemental Fig 3A). Key molecular contacts with side chains were also maintained, such as the “carboxylate clamp” interactions, involving coordination of tauC3’s Asp421 side chain and carboxylate-terminus by cationic Lys30 and Lys95 side chains in CHIP’s TPR domain (Fig 4B). However, we did note a few unique interactions, including between tauC3’s Asp418 and CHIP’s Lys72 (Fig 4C). Critically, this co-structure also suggested why phosphorylation of S416 blocks binding to CHIP. Specifically, we noted that tauC3’s Ser416 was in close proximity with CHIP’s Asp134 (Fig 4D). Thus, phosphorylation of tauC3’s S416 would be expected to create a dramatic interference at this site, involving both steric and electrostatic clashes. To test this idea, we mutated CHIP’s Asp134 (D134A) or the adjacent Phe131 residue (F131A), to alanine and tested binding of these CHIP variants to tauC3 peptides (Fig 4E, Supplemental Fig 3C). Introducing the F131A mutation seemed to damage the integrity of CHIP’s TPR domain because it had a lower melting temperature (Fig 4E), weakened binding to both WT tauC3 and tauC3 S416E peptides (Fig 4E) and poor ubiquitination activity (Supplemental Fig 3B), making this protein a poor tool for further use. Fortunately, installing D134A into CHIP was tolerated; for example, it bound normally to WT tauC3 in DSF experiments (Fig 4E). Importantly, consistent with the design, CHIP D134A even retained significant binding to tauC3 S416E peptide (Fig 4E). To independently verify this finding, we performed FP assays and confirmed that CHIP D134A bound normally to WT tauC3 (K_i_ 5.57 ± 0.34 μM) and that it regained the ability to bind phosphorylated tauC3 peptide (K_i_ 1.84 ± 0.23 μM) (Fig 4F,G). Together, these results support a model in which phosphorylation at S416 weakens binding to CHIP via clashes with D134 in the TPR domain. Interestingly, the D134 residue is highly conserved in evolution (Supplemental Fig 3D,E), suggesting that this residue could be an important “sensor” of phosphorylation.

**Fig 4.**
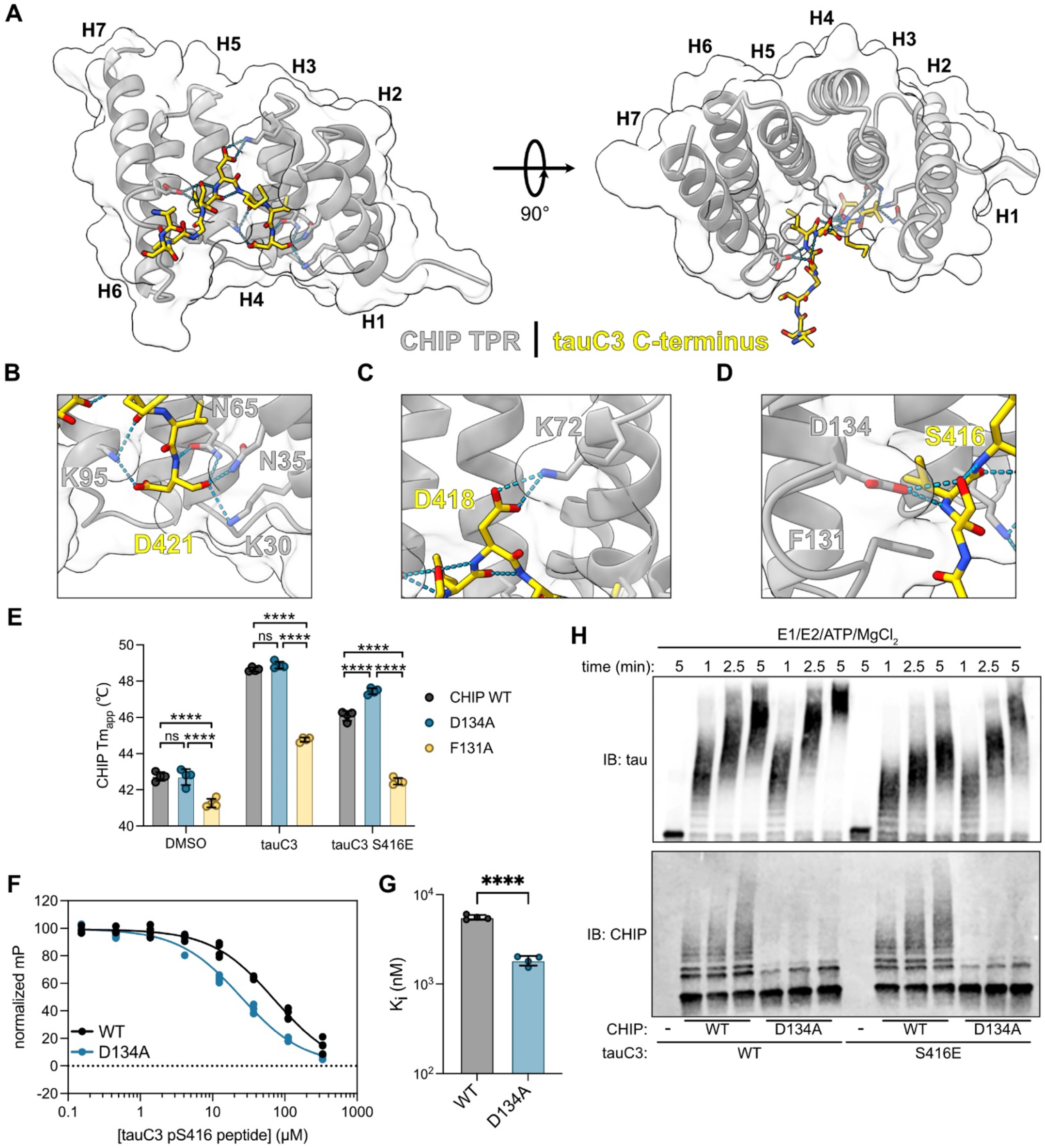
Structural basis for CHIP binding to tauC3 and inhibition by phosphorylation. **(A)** 1.8 Å crystal structure of the CHIP TPR domain bound to a 10-mer tauC3 C-terminal peptide. The CHIP TPR domain is depicted in gray, while the tauC3 peptide is in yellow. Helices 1-7 of the CHIP TPR domain are numbered H1-H7. **(B)** Close-up view of interactions of tauC3 D421 and C-terminus with CHIP carboxylate-clamp residues K95, N65, N35, and K30. **(C)** Close-up view of interaction of tauC3 D418 with CHIP carboxylate-clamp residue K72. **(D)** Close-up view of interactions of tauC3 S416 with CHIP carboxylate-clamp residue D134 and proximity to CHIP F131. **(E)** Apparent melting temperatures (Tm_app_) of CHIP WT or mutants in the absence or presence of 10-mer tau peptides as derived from. Statistical significance was determined by two-way ANOVA with Bonferroni’s post-hoc analysis (ns = not significant, ****p<0.0001, n = 4). **(F)** Competition FP experiment showing displacement of fluorescent tracer from WT or D134A CHIP TPR domain by tauC3 pS416 10-mer peptide. Samples were performed in quadruplicate and normalized to DMSO control. **(G)** Inhibition constants for tauC3 pS416 peptide for WT or D134A CHIP as derived from (F). Statistical significance was determined by unpaired student’s t-test (****p<0.0001, n = 4). **(H)** *In vitro* ubiquitination of tau proteoforms by WT or D134A CHIP. Samples were collected at the denoted timepoints, quenched in SDS-PAGE loading buffer, and analyzed by western blot.

We hypothesized that the CHIP D134A variant might partially restore ubiquitination of pS416 tauC3. To test this idea, we compared the ability of WT or D134A CHIP to ubiquitinate tauC3 or tauC3 S416E *in vitro* (Fig 4H). As expected, we observed more rapid and robust ubiquitination of tauC3 S416E by D134A CHIP, compared to WT CHIP. Interestingly, we also observed that D134A CHIP had relatively more activity against WT tauC3 as well, suggesting that it might be inherently more active. Auto-ubiquitination of CHIP is known to inhibit its function^45^, so we reasoned that D134A might impact this regulatory step. Indeed, we found that, while the D134A mutant CHIP was adept at ubiquitinating substrates, it had significantly reduced autoubiquitination activity (Fig 4H). CHIP is a dimeric protein that assembles into higher order oligomers that retain E3 ubiquitin ligase activity^19^. Thus, the engineered D134A CHIP variant can be used as a tool to study the biological and pathological roles of tauC3, overcoming even the pS416 modification.

### TauC3 co-accumulates with serine 416 phosphorylation in Alzheimer’s disease patient brains

TauC3 is known to accumulate in AD brains and is often used as a pathological biomarker^30^. This finding is somewhat surprising because tauC3 is an excellent substrate for CHIP^36^. Based on our results, we reasoned that this build-up might occur, in part, because tauC3 is phosphorylated at pS416 – preventing its binding to CHIP. To test this idea, we measured whether the tauC3 and pS416 epitopes might co-localize in the brains of AD patients. We performed multiplexed immunofluorescence for tauC3 (C3 antibody, Invitrogen) and tau pS416 (D7U2P, Cell Signaling Technology) on fixed, post-mortem human brain sections of the hippocampal CA1/CA2 and subiculum regions from donors with increasing levels of AD pathology (Fig 5A, Supplemental Fig 4A). Using this approach, we observed significant increases in both tau pS416 and tauC3 across disease progression (Fig 5B, 5C) and the appearance of the two PTMs strongly correlated by Pearson’s analysis (r = 0.7681, p = 0.0035; Fig 5D). In the soma of dystrophic neurons, we often observed overlapping pathology for tauC3 and tau pS416 (Fig 5A – inset), suggesting these two tau PTMs could be contributing to disease when co-occurring. These results support the model that Ser416 phosphorylation stabilizes tauC3, contributing to its accumulation *in vivo*.

**Fig 5.**
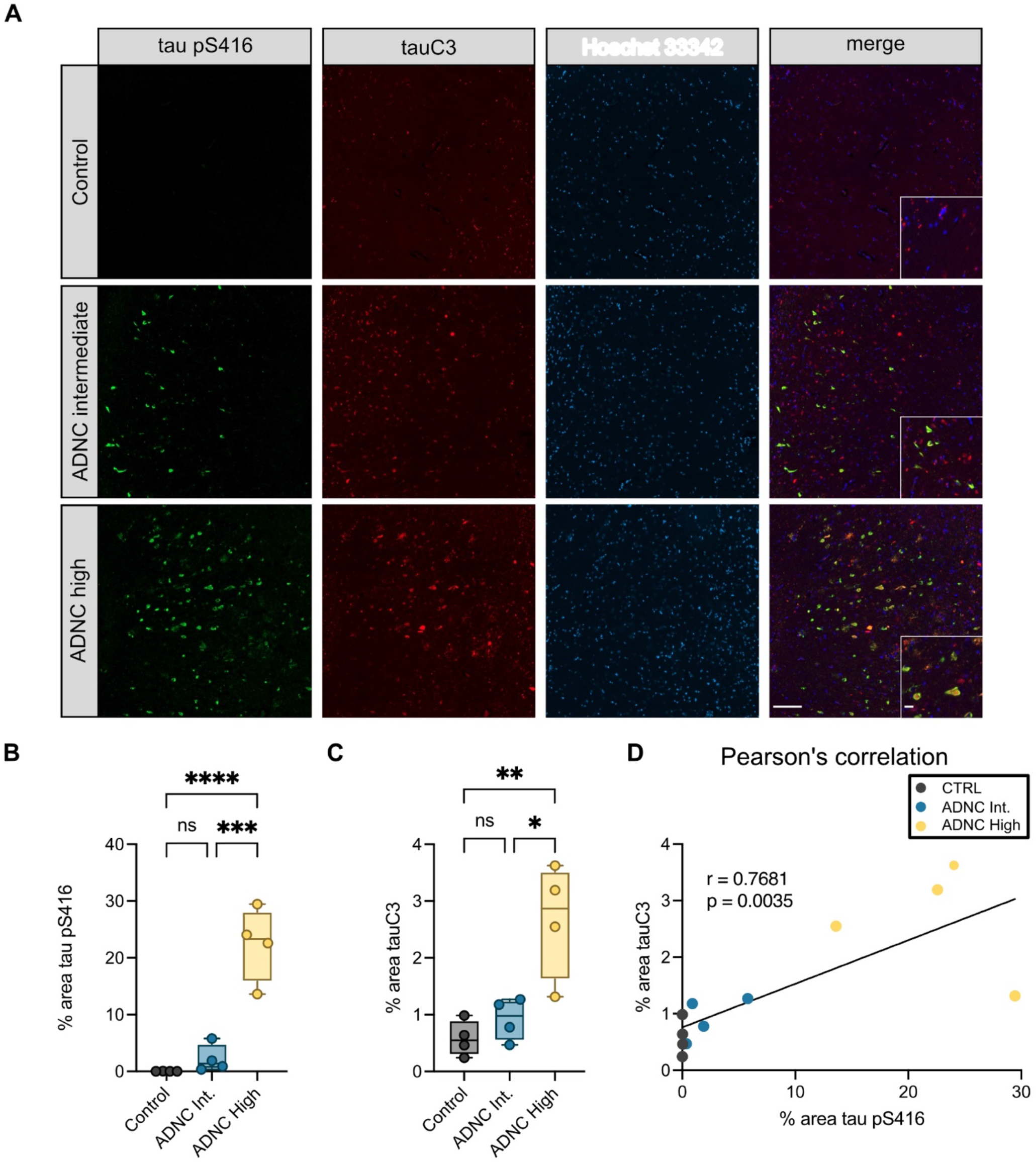
TauC3 co-accumulates with Serine 416 phosphorylation in Alzheimer’s disease patient brains. **(A)** Representative micrographs of immunofluorescence staining of tau pS416 (green), tauC3 (red), and nuclei (Hoechst 33342, blue) from the hippocampal CA1/CA2 region of human patient brains across increasing spectrum of Alzheimer’s Disease Neuropathological Change (ADNC) scoring. Scale bars = 50 μM; insets = 10 μM. **(B)** Quantification of tau pS416 staining across ADNC score. Analysis was performed on four patient samples for each score. Statistical significance was determined by one-way ANOVA with Tukey’s post-hoc analysis (ns = not significant, ***p<0.001, ****p<0.0001). **(C)** Quantification of tauC3 staining across ADNC score. Analysis was performed on four patient samples for each score. Statistical significance was determined by one-way ANOVA with Tukey’s post-hoc analysis (ns = not significant, ***p<0.001, ****p<0.0001). **(D)** Pearson’s analysis showing correlation of increasing tau pS416 and tauC3 across ADNC score.

### MARK2 inhibits CHIP-dependent ubiquitination of tauC3 by phosphorylating serine 416

Many tau kinases have been identified^46^ and we wondered which of these kinase(s) might phosphorylate tauC3 Ser416. Previous studies have implicated Ca^2+^-calmodulin-dependent protein kinase II (CaMKII) in the phosphorylation of Ser416 on FL tau^47,48^. However, the relevant CaMKII isoforms are specific to the central nervous system^49^, and are not expressed in the HEK293 or *Sf9* cells that we employed. Thus, we turned our attention to the CaMKII-like kinase microtubule affinity regulating kinase 2 (MARK2/Par-1) as a likely candidate, because it has also been shown to phosphorylate Ser416 in FL tau^39^. As a first step, we incubated FL tau, tauC3, or phosphomimetic tauC3 S416E with recombinant MARK2 *in vitro* and probed for Ser416 phosphorylation by western blot. Indeed, we observed robust phosphorylation of Ser416 in both FL and tauC3, as well as slowed in-gel mobility that is consistent with tau phosphorylation (Fig 6A). In ubiquitination assays, the rate of ubiquitination for tauC3 was slowed following phosphorylation by MARK2 (Fig 6B-D). Together, these data suggest that MARK2-mediated phosphorylation of tauC3 at S416 could be one of the contributing factors to the inhibition of CHIP-mediated ubiquitination, with important implications for the accumulation of this tau proteoform in disease.

**Fig 6.**
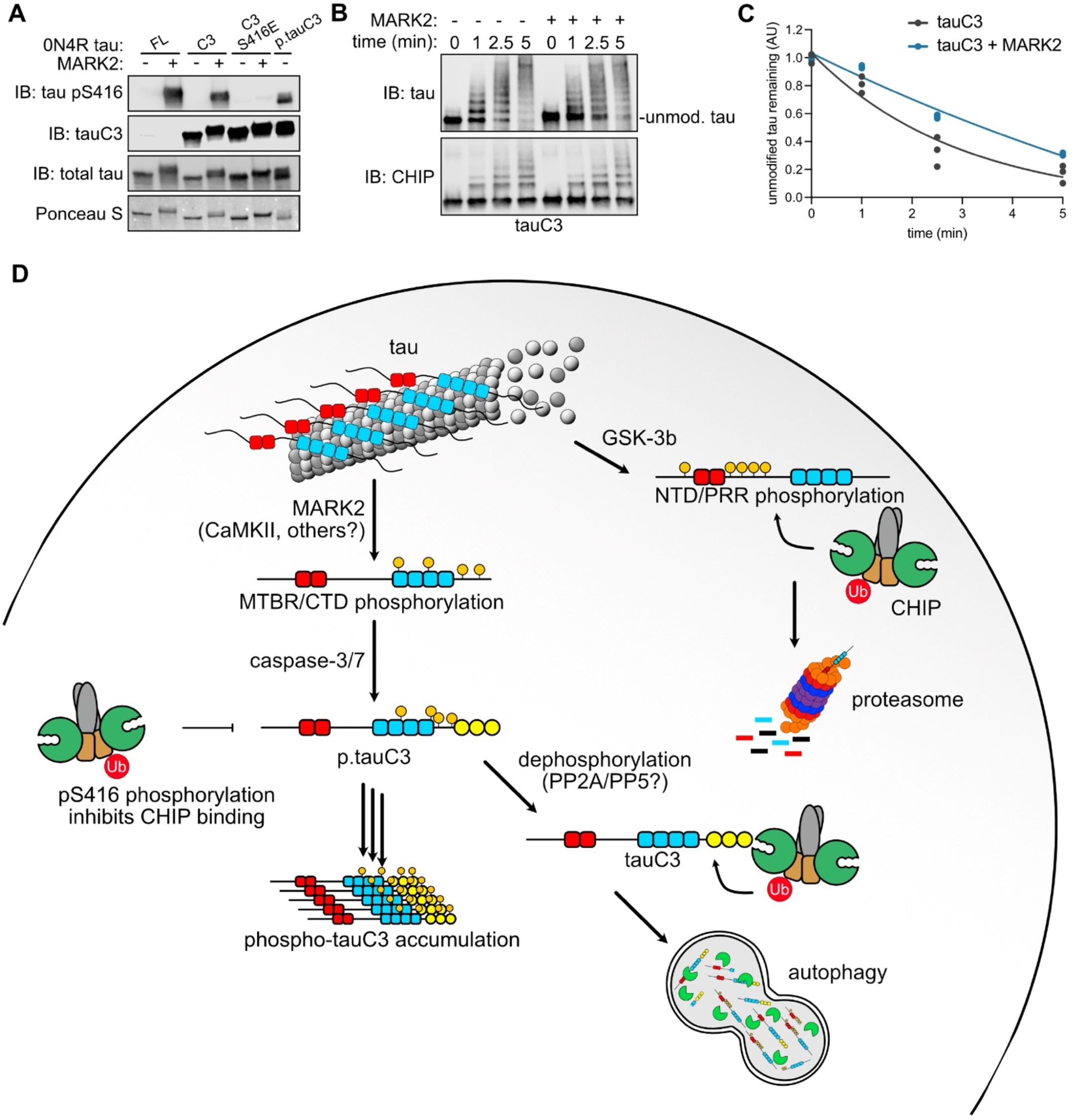
MARK2 inhibits CHIP-dependent ubiquitination of tauC3 by phosphorylating serine 416. **(A)** *In vitro* phosphorylation of varying tau proteoforms by MARK2. Varying tau species were incubated overnight with recombinant MARK2, and relative phosphorylation of Ser416 was quantified by western blot. Ponceau S was used as loading control. **(B)** *In vitro* ubiquitination of unmodified tauC3 or MARK2-phosphorylated tauC3 by CHIP. Samples were collected at the denoted timepoints, quenched in SDS-PAGE loading buffer, and analyzed by western blot. **(C)** Quantification of unmodified tau remaining in (C). Samples were normalized to 0-minute time point and curves were fit using one-phase exponential decay (n = 3). **(D)** Cartoon depicting the role of tau PTMs on interactions with CHIP and shuttling to various degradation pathways.

## DISCUSSION

Mass spectrometry studies on tau isolated from the brains of AD patients has shown that this protein is subject to extensive PTMs, including phosphorylation, proteolysis, ubiquitination, and acetylation^22,50,51^. Many of these modifications correlate with disease progression and they are being pursued as promising, diagnostic biomarkers. Yet, the molecular mechanisms connecting these PTMs to tau proteostasis are often lacking^22,52 53^. For example, tau proteoforms that are phosphorylated at pT181 and pT217 are strong biomarkers for AD^54^, but it is not clear whether these phosphorylation events are directly impacting tau accumulation. In other words, are they a cause or consequence? Here, we focus on understanding the intersection between two tau PTMs that are strongly linked to AD: phosphorylation at S416 and caspase cleavage to produce tauC3. We considered this system to be a good model for asking questions about the mechanistic roles of tau phosphorylation because recent work has clearly identified the binding site of the E3 ligase CHIP on tauC3^36^.

Using *in vitro* and cellular assays, we find that MARK2-mediated phosphorylation of tauC3 at pS416 is sufficient to block binding to CHIP, resulting in its enhanced accumulation in cells. Moreover, the appearance of pS416 and tauC3 are coincident in AD patient samples, supporting a role of a single phosphorylation in affecting tau proteostasis. Based on these collective findings and in concert with observations from the literature, we propose a model to explain the hierarchy of these tau PTMs (Fig 6D). In this speculative model, FL tau can either be phosphorylated by MARK2/CaMKII, to generate phosphosites in the MTBRs^39^, or GSK-3β, which generates sites in the NTDs and PRR^26^. If the GSK3β pathway predominates, then those tau proteoforms seem likely to bind directly to CHIP and are cleared through the UPS^26^. This result is broadly consistent with our observations that okadaic acid treatment enhances binding of FL tau to CHIP (see Fig 2). However, if tau is proteolyzed by caspase-3/7 to generate tauC3, then this protein binds directly to CHIP through its new, C-terminal EEVD-like motif^36^ and it is then degraded by the lysosomal-autophagy pathway^55^. Caspases are known to be active during in adult neurons in a non-apoptotic signaling pathway^30^, so it seems likely that tauC3 might be normally produced to feed into this degradation pathway. Here, we show that phosphorylation of tauC3 at pS416 is sufficient to block degradation, creating an intersection of two tau PTMs: proteolysis and phosphorylation. We propose that this event creates a requirement for dephosphorylation, likely by PP2A or PP5^56,57^, the major enzymes that use tau as a substrate. It is not clear whether MARK2/CaMKII activity occurs before or after caspase cleavage. However, phosphorylation of tau at a nearby residue, S422, blocks caspase cleavage^58^ and, more generally, phosphorylated tau is a poor substrate for caspases^59^, which could be an important determinant of this hierarchy. While this proposed model is likely to be a simplistic representation, it does serve to highlight how CHIP plays multiple, but non-overlapping, roles in tau proteostasis. For example, CHIP binds different tau proteoforms (*e*.*g*. tauC3, FL tau) in distinct ways and tau phosphorylation (*e*.*g*. GSK3β-mediated, pS416) can have diametrically opposite effects on CHIP’s apparent affinity. Moreover, even the major degradation pathway (*e*.*g*. proteasome vs. lysosome) used by CHIP when pairing with each proteoform can be different. When properly balanced, we envision that CHIP uses both the GSK3β and MARK2/CaMKII pathways, along with the canonical Hsp70-mediated mechanism^5,19^, to remove a wide variety of tau proteoforms (Fig 6D). This broad recognition activity might explain why CHIP is such a strong regulator of tau^6^, despite the fact that tau exists in so many different proteoforms. However, an over-production of tauC3, combined with aberrant phosphorylation of pS416, might give rise to a tendency to form NFTs under specific conditions, especially if CHIP is limiting^36^. In theory, disease-associated dysregulation might occur at any of the proposed steps, such as activation of caspase 3/7 and/or MARK2 or a decrease in the flux through the degradation pathways as a function of aging.

One of the goals of understanding the molecular mechanisms of tau PTMs is to identify putative drug targets and link these to specific tau proteoform biomarkers^60^. Our studies showed that MARK2 phosphorylation of tauC3 slows CHIP-dependent ubiquitination (see Fig 6), suggesting that MARK2 inhibitors might be beneficial, while also focusing on tauC3 as the key substrate. MARK2 had previously been linked to AD; for example, MARK2-mediated phosphorylation weakens the interaction of tau with microtubules and promotes tau’s cytosolic accumulation^39,61^ and certain MARK2 sites, such as Ser262, Ser324, and Ser396, are elevated in AD^62^. Moreover, MARK2 phosphorylation has been shown to promote tau liquid-liquid phase separation, a process that might promote the transition into amyloid fibrils^63^. Our results add to this series of observations, suggesting that activity at pS416 might be particularly important for regulating tauC3.

## MATERIALS AND METHODS

### Cell lines and culture conditions

HEK 293T cells (ATCC) were cultured in Dulbecco’s Modified Eagle Medium (DMEM) supplemented with 10% Fetal Bovine Serum (FBS) and penicillin/streptomycin. Parent HEK 293 Flp-In T-REx cells (Thermo Fisher) were cultured in 10% FBS-DMEM supplemented with 1X penicillin/streptomycin, 15 μg/mL blasticidin, and 100 μg/mL zeocin. All mammalian cells were maintained at 37 °C / 5% CO_2_ in a humidified incubator.

For construction of the stable NanoBiT reporter cell line, parent HEK 293 Flp-In T-REx cells were harvested by trypsinization, resuspended in electroporation buffer containing 5 μg eGFP-tau plasmid DNA and 45 μg pOG44 plasmid DNA. Cells were electroporated using program Q-001 on a Lonza 4D-Nucleofector. Electroporated cells were immediately diluted in 10% FBS-DMEM and cultured for 48 hours to recover. Following recovery, cells were selected with 100 μg/mL hygromycin until single colonies arose. Colonies were subsequently picked and screened for dox-inducible expression of eGFP-tau by fluorescence microscopy (Echo Revolve) and western blotting.

### NanoBiT live cell PPI assay

HEK293T cells were seeded in poly-D-lysine coated 96-well plates (Corning, flat bottom, white opaque) in Opti-MEM. Cells were transfected with 50 ng each SmBiT/LgBiT DNA using Lipofectamine 3000 (Thermo Fisher) in Opti-MEM according to the manufacturer’s instructions. Transfections were performed for 24 hours, after which the media was replaced with fresh Opti-MEM containing DMSO or 30 nM okadaic acid. Cells were incubated for an additional 16 hours at 37 °C / 5% CO_2_ in a humidified incubator. Following this incubation, 12.5 uL 1X Nano-Glo live cell luciferase reagent (Promega) diluted in LCS buffer (Promega) was added to each well, and samples were incubated for 10 minutes at room temperature in the dark to allow luminescence levels to stabilize. Luminescence recordings were performed on a SpectraMax M5 plate reader with 500 ms integration time in well-scan mode acquiring 9 readings per well. Un-transfected cells were used to obtain background measurements which were subtracted from observed luminescence values.

### Protein purification

Recombinant human CHIP was expressed from a pMCSG7 construct with N-terminal tobacco etch virus (TEV) – cleavable 6His-tag. pMCSG7-CHIP was transformed into BL21DE3 (New England Biolabs) *E. coli* and grown in terrific broth (TB) to OD600 = 0.5 at 37 °C. Cells were cooled to 16 °C, induced with 500 μM isopropyl β-D-1-thiogalactopyranoside (IPTG), and grown overnight. Cells were collected by centrifugation, resuspended in binding buffer (50 mM Tris pH 8.0, 10 mM imidazole, 500 mM NaCl) supplemented with cOmplete protease inhibitors (Roche), and sonicated. The resulting lysate was clarified by centrifugation and the supernatant was applied to Ni^2+^-NTA His-Bind Resin (Novagen). Resin was washed with binding buffer and His wash buffer (50 mM Tris pH 8.0, 30 mM imidazole, 300 mM NaCl), and then eluted from the resin in His elution buffer (50 mM Tris pH 8.0, 300 mM imidazole, 300 mM NaCl). Following this step, the N-terminal His tag was cleaved by overnight dialysis with TEV protease at 4 °C and purified by size exclusion chromatography (SEC) (HiLoad Superdex-200 16/600 column, GE Healthcare) in CHIP storage buffer (50 mM HEPES, 10 mM NaCl, pH 7.4).

CHIP TPR Domain (human, AA 22–154) was expressed from a pMCSG7 construct with an N-terminal TEV-cleavable 6xHis tag, as previously described^36^. *E. coli* were grown in TB at 37 °C, induced with 1 mM IPTG in log phase, cooled to 16° C and grown overnight. Ni-NTA purification and tag removal were conducted as for full-length CHIP. Protein was further purified on a Mono S cation exchange column (GE Healthcare) and stored in CHIP TPR storage buffer (10 mM Tris, 150 mM NaCl, 2 mM DTT, pH 8.0).

Recombinant, unmodified 0N4R tau proteins were expressed from a pMCSG7 construct with N-terminal TEV – cleavable 6His-tag. pMCSG7-tau was transformed into BL21DE3 *E. coli* and grown in terrific broth (TB) to OD600 = 0.5 at 37 °C. Betaine (10 mM) and NaCl (500 mM) were added to media and tau was induced by addition of IPTG (200 μM) for 3 hours at 30 °C. Cells were collected by centrifugation, resuspended in tau lysis buffer (1X D-PBS, 2 mM MgCl_2_, 1 mM DTT, 1 mM EDTA, pH 7.4) supplemented with cOmplete protease inhibitor and 1 mM phenylmethylsulfonyl fluoride (PMSF), and lysed by sonication. The lysate was clarified by centrifugation and applied to cOmplete His-tag purification resin (Sigma Aldrich) for 1 hour at 4 °C with rotation. The resin was washed with tau lysis buffer and bound protein was eluted with tau elution buffer (1X D-PBS, 300 mM imidazole, 200 mM NaCl, 5 mM β-mercaptoethanol, pH 7.4). His-tags were removed by addition of TEV protease and overnight dialysis into tau buffer (1X D-PBS, 2 mM MgCl_2_, 1mM DTT, pH 7.4) at 4 °C. Proteins were subsequently concentrated and purified by reverse-phase HPLC as previously described^64^. Solvent was removed by lyophilization, and the resulting protein samples were resuspended in tau buffer.

Phosphorylated tauC3 was expressed in Sf9 insect cells from a 438B pFastBac vector containing an engineered 6xHis tag and a TEV protease cleavage site (Addgene). Following two rounds of virus amplification, the protein was expressed in Sf9 cells by infecting with recombinant baculovirus at 20 to 40 μL virus per million cells in culture flasks and incubated for 3 days at 27 °C with shaking at 120 rpm. Cells are then collected by spinning at 1500 rpm for 20 minutes at 4 °C and stored at -80 °C. The cells are resuspended in Ni lysis buffer (20 mM Tris, pH 8.0, 500 mM KCl, 10 mM imidazole, 10% glycerol) supplemented with EDTA-free protease inhibitor cocktail (Roche) and 6 mM beta-mercaptoethanol. Cells were lysed in a Dounce homogenizer, and the lysates were boiled for 20 mins to denature and precipitate nearly all proteins except for tau. The lysate was centrifuged (40000 rpm) at 4 °C for 30 min to clarify and the supernatant was incubated with HisPur Ni-NTA resin (Thermo Scientific) at 4 °C for 1 hour. The resins were then washed and eluted with elution buffer (20 mM Tris-HCl, 100 mM KCl, 6 mM β-mercaptoethanol, 300 mM imidazole, pH 8.0). The fractions containing tau, as judged by SDS-PAGE, were dialyzed with a dialysis buffer (50 mM Tris-HCl, 100 mM KCl, 6 mM β-mercaptoethanol, 5% glycerol, pH 8.0) containing TEV protease to cleave the His tag at 4 °C overnight. The dialyzed proteins were concentrated and purified by SEC (HiLoad Superdex-200 16/600 column, GE Healthcare) into tau buffer. Phosphorylation was confirmed by gel-shift and western blotting with phospho-specific antibodies.

MARK2 phosphorylation of recombinant tau proteins was performed as previously described^39^ with slight modification. In brief, recombinant GST-tagged MARK2 (Promega) was incubated with tau at 30 °C at 1:100 ratio of MARK2:tau in phosphorylation buffer (25 mM PIPES, 100 mM NaCl, 5 mM MgCl_2_, 2 mM EGTA, 1 mM DTT, 1 mM benzamidine, 0.5 mM PMSF, 1 mM ATP, pH 6.8) for 18 hours. Following, samples were incubated with equilibrated Pierce glutathione resin (Thermo Fisher) for 1 hour to remove MARK2, followed by desalting and buffer exchange into tau buffer over Zeba protein desalting columns (Thermo Fisher).

### Peptide synthesis

Peptides were synthesized by Fmoc solid phase peptide synthesis on a Syro II peptide synthesizer (Biotage) at ambient temperature and atmosphere on a 12.5 μmol scale using pre-loaded Wang resin. Coupling reactions were conducted with 4.9 eq of HCTU (O-(1H-6-chlorobenzotriazole-1-yl)-1,1,3,3-tetramethyluronium hexafluoro-phosphate), 5 eq of Fmoc-AA-OH and 20 eq of N-methylmorpholine (NMM) in 500 μL of N,N dimethyl formamide (DMF). Reactions were run for 8 min while shaking. Each position was double coupled. Fmoc deprotection was conducted with 500 μL 40% 4-methypiperadine in DMF for 3 minutes, followed by 500 μL 20% 4-methypiperadine in DMF for 10 minutes, and six washes with 500 μL of DMF for three minutes. Acetylation was achieved by reaction with 20 eq acetic anhydride and 20 eq NMM in 500 μL DMF for 1 h while shaking. Peptides were cleaved with 500 μL of cleavage solution (95% trifluoroacetic acid 2.5% Water 2.5% triisopropylsilane) while shaking for 1 h. Crudes were precipitated in 10 mL cold 1:1 diethyl ether : hexanes. Peptide crudes were solubilized in a 1:1:1 mixture DMSO: water: acetonitrile and purified by HPLC on an Agilent Pursuit 5 C18 column (5 mm bead size, 150 mm × 21.2 mm) using an Agilent PrepStar 218 series preparative HPLC. The mobile phase consisted of A: Water 0.1% Trifluoroacetic acid and B: Acetonitrile 0.1% Trifluoroacetic acetic acid. Solvent was removed under reduced atmosphere and 10 mM DMSO stocks were made based on the gross peptide mass. Purity was confirmed by LC/MS. Stocks were stored at −20 °C. Fluorescence polarization tracer peptides and tau phospho-peptides were synthesized by Genscript.

### Western blotting

Prepared samples were separated on precast 4-20% SDS-PAGE gradient gels (Bio-Rad) for 35 minutes at 200V. Proteins were transferred to nitrocellulose membranes using a Trans-Blot Turbo Transfer System (Bio-Rad). Membranes were blocked in Odyssey TBS Blocking Buffer (Licor) for 1 hour at room temperature and then incubated in primary antibody in 1X Tris-Buffer saline containing 0.05% Tween-20 (1X TBS-T), plus 5% non-fat dry milk powder, and 0.02% NaN_3_ overnight at 4 °C with rotation. The following day, membranes were washed 3 times in 1X TBS-T and then incubated in secondary antibody diluted 1:10000 in Odyssey TBS Antibody Diluent (Licor) for one hour at room temperature. Following incubation with secondary antibody, membranes were washed 3 times in 1X TBS-T and imaged on an Odyssey Fc Imaging System (Licor). Quantification was performed by densitometry analysis in ImageJ (NIH).

### CHIP-tau binding ELISA

Purified CHIP (1 μM) or buffer-matched control was immobilized in 96-well plates (Fisher Scientific, non-sterile, clear, flat-bottom) in CHIP buffer (50 mM HEPES, 10 mM NaCl, pH = 7.4) overnight at 37 °C. The protein sample was removed, and wells were washed 3X with phosphate-buffered saline with 0.05% Tween-20 (PBS-T) for 3 minutes with rotation at room temperature. Tau samples were prepared as a 3-fold dilution series in tau binding buffer (25 mM HEPES, 40 mM KCl, 8 mM MgCl2, 100 mM NaCl, 0.01% Tween, 1 mM DTT, pH 7.4) and incubated at RT for 3 hours with rotation. Tau was removed, and wells were washed as described. Samples were blocked in 5 % non-fat dry milk in tris-buffered saline (TBS), and then incubated with primary anti-tau (Santa Cruz Biotech, 1:2000) followed by HRP-conjugated secondary antibody (Anaspec, 1:2000). Antibodies were dissolved in 1X TBS with 0.05% Tween-20. Incubations were performed for 1 hour at RT with rotation, separated by wash steps, as described^23^. TMB substrate (Thermo Fisher) was then added to the wells and incubated for 15 min at RT, followed by quenching with 1 M HCl. Absorbance readings were performed on a SpectraMax M5 plate reader at OD_450_. The data was background subtracted to buffer only controls, normalized to maximal binding, and binding curves were fit using non-linear regression in Prism 9.0 (GraphPad).

### In vitro ubiquitination assays

In preparation for *in vitro* ubiquitination reactions, four 4× stock solutions were prepared containing (1) Ube1 + UbcH5c (400 nM Ube1 and 4 μM UbcH5c), (2) Ubiquitin (1 mM Ub), (3) CHIP + substrate (4 μM CHIP and 4 μM substrate) and (4) ATP + MgCl_2_ (10 mM ATP and 10 mM MgCl_2_) in ubiquitination assay buffer (50 mM Tris pH 8.0, 50 mM KCl). Ubiquitination reactions were generated by adding 10 μL of each 4× stock, in order from 1 to 4, for a final volume of 40 μL (100 nM Ube1, 1 μM UbcH5c, 250 μM ubiquitin, 2.5 mM ATP, 2.5 mM MgCl_2_, 1 μM CHIP and 1 μM substrate). Reactions were then incubated at room temperature, and 10 μL aliquots were collected at each time point and quenched in 5 μL 3× SDS–PAGE loading buffer. Samples were separated by SDS-PAGE and analyzed by western blotting. Quantification of unmodified tau remaining was performed by densitometry analysis in ImageJ (NIH).

### Mapping of tau PTMs by liquid chromatography with tandem mass spectrometry

A sample of protein (8 μg) was denatured and reduced with 8 M urea in 100 mM ammonium bicarbonate (pH 8) buffer and 100 mM dithiothreitol (DTT) at 60 °C for 30 minutes, followed by alkylation with 100 mM iodoacetamide at room temperature in the dark for 1 hour. The sample was then incubated 4 hours with trypsin (1:20 weight/weight) at 37 °C. The peptides formed from the digestion were further purified by C18 ZipTips (Millipore) and analyzed by LC-MS/MS. The MS/MS analyses were conducted using either an Q Exactive Plus Orbitrap (QE) or a Fusion Lumos Orbitrap (Lumos) mass spectrometer (Thermo Scientific). Higher-energy collisional dissociation was used to produce fragmented peptides. The mass resolution of precursor ions was 70000 on the QE and 120000 on the Lumos. The mass resolution of fragment ions was 17500 on the QE and 30000 on the Lumos, respectively. The LC separation was carried out on a NanoAcquity UPLC system (Waters) for both the QE and the Lumos. The LC linear gradient on the QE was increased from 2 -25% B (0.1% formic acid in acetonitrile) over 48 mins followed by 25 -37% B over 6 mins and then 37 – 40% B over 3 mins at a flow rate of 400 nL/min. The LC linear gradient on the Lumos was increased from 2 - 5% B over 3 mins followed by 5 - 30% B over 72 mins and then 30 – 50% B over 2 mins at a flow rate of 300 nL/min. The acquired MS/MS raw data was converted into peak lists using an in-house software PAVA and then analyzed using Protein Prospector search engine. The Max. missed cleavages was set to 2. The precursor / fragment mass tolerances were set at 20 ppm / 20 ppm for the QE and 10 ppm / 20 ppm for the Lumos. For all peptides, phosphorylation modification at serine, threonine, and tyrosine residues was selected. Visualization of tau PTMs was performed using Protter^65^.

### Differential scanning fluorimetry

DSF was performed with a 10 μL assay volume in 384-well Axygen quantitative PCR plates (Fisher Sci) on a qTower^3^ real-time PCR thermal cycler (Analytik Jena). Fluorescence intensity readings were taken over 70 cycles in “up-down” mode, where reactions were heated to desired temp and then cooled to 25 °C before reading. Temperature was increased 1 °C per cycle. Each well contained 5 μM CHIP, 5× Sypro Orange dye (Thermo Fisher), and varying concentrations of peptide in DSF assay buffer (25 mM HEPES pH 7.4, 50 mM KCl, 1 mM TCEP, 0.2% CHAPS, 1%DMSO). Fluorescence intensity data was truncated between 30-60 °C, plotted relative to temperature, and fit to a Boltzmann Sigmoid in GraphPad Prism 9.0. CHIP apparent melting temp (Tm_app_) was calculated based on the following equation:

Y=Bottom+((Top-Bottom)/(1+exp(*Tm*-*T*/Slope)))

### Fluorescence polarization

Fluorescence polarization (FP) assays were performed in 18 μL in a Corning black 384 well round bottom low volume plate and measurements made on a SpectraMax M5 multimode plate reader at 21 °C. A 2× stock of CHIP + Tracer was made in CHIP FP assay buffer, so that the final assay concentration of CHIP was 1.58 μM and tracer was 20 nM. The 2× peptide competitor stocks were prepared in CHIP FP Dilution buffer (25 mM HEPES pH 7.4, 50 mM KCl 0.01% Triton X-100, 2% DMSO) in three-fold dilutions. 2× CHIP + Tracer and peptide competitor solutions were mixed at equal volumes and incubated at 25°C in the dark for 15 minutes. Raw polarization (mP) values were background subtracted to tracer alone and plotted relative to log_10_ (competitor). Data was fit to the model for [inhibitor] versus response (three parameters) in Graphpad Prism 9.0. IC_50_ values were calculated based on the equation:

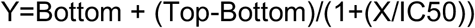

K_i_ values were calculated as previously described^66^ using the equation:

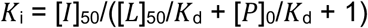

### Co-immunoprecipitation assays

Cells were washed once in 1X Dulbecco’s Phosphate Buffered Saline (D-PBS) and lysed in ice cold Pierce Immunoprecipitation (IP) lysis buffer (25 mM Tris-HCl pH 7.4, 150 mM NaCl, 1 mM EDTA, 1% NP-40). Lysates were collected by scraping, transferred to 1.5 mL microcentrifuge tubes, and incubated on ice for 15 minutes. Following, lysates were clarified centrifugation at 15,000 RPM for 10 minutes at 4 °C and supernatants were transferred to new microcentrifuge tubes. Relative protein concentrations were determined by bicinchoninic acid assay (Pierce), and samples were normalized to the lowest protein concentration. A representative input sample of 40 μL was collected, mixed with 20 μL 3X SDS-PAGE loading buffer (188 mM Tris-HCl pH 6.8, 3% SDS, 30% glycerol, 0.01% bromophenol blue, 15% β-mercaptoethanol), denatured at 95 °C for 5 minutes, and stored at 4 °C for later analysis.

For tau immunoprecipitations, an aliquot (30 μL) of Protein A/G PLUS-agarose (Santa Cruz Biotech) per condition was collected and washed twice in ice cold IP lysis buffer. Then, an aliquot (30 μL) of resuspended beads and anti-Tau-5 antibody (Thermo Fisher, 1:250) were added to prepared lysates and incubated at 4 °C overnight with rotation. The following day, beads were collected by centrifugation and supernatants were removed. Beads were washed 3 times in ice cold IP lysis buffer, and bound proteins were eluted by addition of 60 μL 1X SDS-PAGE loading buffer and denaturing at 95 °C for 5 minutes. Inputs and eluates were subsequently analyzed by SDS-PAGE followed by western blotting. Quantification was performed by densitometry analysis in ImageJ (NIH).

### qPCR

Relative quantitation of *MAPT* gene expression was performed as previously described^67^. Briefly, cells were collected by centrifugation and RNA was extracted via Quick-RNA Miniprep Kit (Zymo Research) and complementary DNA (cDNA) was synthesized via SensiFast cDNA Synthesis Kit (Meridian Bioscience), both according to the manufacturer’s instructions. Samples were prepared for qPCR in both technical and biological triplicates in 5-μl final volumes using SensiFAST SYBR Lo-ROX 2X Master Mix (Meridian Bioscience) with qPCR primers (Integrated DNA Technologies) at a final concentration of 0.2 μM and cDNA diluted at 1:3. qPCR was performed on a QuantStudio 6 Pro Real-Time PCR System (Applied Biosystems) using QuantStudio Real Time PCR software (v.1.3) with the following Fast 2-Step protocol: (1) 95 °C for 20 s; (2) 95 °C for 5 s (denaturation); (3) 60 °C for 20 s (annealing/extension); (4) repeat steps 2 and 3 for a total of 40 cycles; (5) 95 °C for 1 s; (6) ramp 1.92 °C s^−1^ from 60 °C to 95 °C to establish melting curve. Expression fold changes were calculated using the ∆∆Ct method and normalized to housekeeping gene *ACTB*.

### X-ray crystallography

The protein solution was prepared by mixing a 1:2 molar ratio of human CHIP-TPR, at 6 mg/ml, and the 6-mer or 10-mer tau peptide in protein buffer (10 mM Tris-HCl pH 8.0, 150 mM NaCl and 2 mM DTT), and incubated on ice for 30 min. Crystals of the complex were grown at room temperature by hanging-drop by mixing 100 nL of the protein solution with 100 μL of the crystallization condition (0.1 M CaCl_2_, 0.1 M HEPES (pH 7.4), 28% PEG 4K (10mer) or 25% PEG 3350 (6mer)) by Mosquito Nanoliter Dropsetter (TTPLabtech). Crystals appeared within 48 h and were harvested ∼ 1 week after setup by flash-freezing in liquid nitrogen using a cryogenic solution of 50% MPD in the crystallization condition. Data were collected at Lawrence Berkeley National Laboratory Advanced Light Source beamline 8.3.1. Diffraction images were processed using Xia2 with the Dials pipeline^68^. Automatic molecular replacement was performed using the online Balbes tool^69^. The resulting structure models were refined over multiple rounds of restrained refinement and isotropic B-factor minimization with Phenix^70^. Structural visualization and rendering for figures were performed using UCSF ChimeraX-1.4. See Supplementary Methods for additional information.

### Histopathology

The de-identified, post-mortem tissues were sourced from the Neurodegenerative Disease Brain Bank at University of California at San Francisco (UCSF; https://memory.ucsf.edu/neurodegenerative-disease-brain-bank). Samples for immunofluorescence were formalin fixed paraffin embedded and cut at 8μm. Slides were deparaffinized and subjected to hydrolytic autoclaving at 121 °C for 10 min in citrate buffer (Sigma, C9999). Following blocking with 10% normal goat serum (Vector laboratories, S-1000), sections were incubated with primary antibodies, tau pS416 (Cell Signaling, 15013) 1:200 and tauC3 (Millipore, MAB5430) 1:250, overnight at room temp. After washing, sections were incubated in secondary antibodies Alexa Fluor goat anti rabbit 488 (Thermo Fisher A11008) and Alexa Fluor goat anti mouse 647 (Thermo Fisher A21235) both 1:500 for 120 minutes at room temp. Sections were then washed and incubated in Hoechst (Life Technologies H3570) 1:5000 for 10 minutes and then rinsed with DI water and coverslipped using Permafluor aqueous mounting medium (Thermo Scientific, TA030FM). Slides were imaged using the Zeiss AxioScan.Z1. Digital images were analyzed using the Zeiss Zen 3.5 (blue edition) Analysis software. To quantify phospho-tau (Ser416) and cleaved tau (tauC3) neuropathology, a pixel intensity threshold was determined using a slide from a clinically diagnosed AD patient with high neuropathologic change and was then applied to all slides. Regions of interest were drawn in the subiculum and the CA1/CA2 of the hippocampus, and the percent area of pixels positive for staining in each region was determined.

## Acknowledgements

This work was supported by grants from the Alzheimer’s Association (to C.N.) and the NIH (AG077842 to M.D.C. and AG068125 to D.R.S, C.S.C. and J.E.G.). Additional support was provided by the BrightFocus Foundation and the Tau Consortium. Human tissue samples were provided by the Neurodegenerative Disease Brain Bank at the University of California, San Francisco, which receives funding support from NIH grants P01AG019724 and P50AG023501, the Consortium for Frontotemporal Dementia Research, and the Tau Consortium.

## Author Contributions

C.M.N and J.E.G. designed the studies and wrote the manuscript. All authors edited the manuscript. C.M.N.,

A.C.T. and M.D.C. conducted biochemical and cell-based experiments, performed data analysis, and generated necessary reagents. K.B. and K.W. performed the crystallography experiments, including data collection and data processing. A.O. performed the histopathology studies. J.E.G., D.A.M., C.S.C. and D.R.S. provided project oversite, data interpretation and funding.

## Additional Information

Supplementary Information is provided, including crystallography information and Supplementary Figures 1-4.

