## Supplemental Information for "Phosphorylation of a Cleaved Tau Proteoform at a Single Residue Inhibits Binding to the E3 Ubiquitin Ligase, CHIP"

**Supplemental Methods:** Details of the crystallography experiments

**Supplemental Fig. 1** Expression of tauC3 in insect cells yields protein that is phosphorylated at multiple sites, including pathogenic residues.
**Supplemental Fig. 2** Pseudo-phosphorylation at serine 416 is sufficient to inhibit CHIP binding.
**Supplemental Fig. 3** Mutation of a conserved residue in CHIP (CHIPD134A) partially restores activity on tauC3 pS416.
**Supplemental Fig. 4** Additional examples of pS416 accumulation in AD and co-localization with tauC3.

**SUPPLEMENTAL METHODS**

| | CHIP-TPR 10mer tau (r26) |
|---|---|
| Wavelength | |
| Resolution range | 61.33 - 1.848 (1.914 - 1.848) |
| Space group | C 1 2 1 |
| Unit cell | 82.34 46.0351 70.7818 90 119.95 90 |
| Total reflections | 112271 (11324) |
| Unique reflections | 19695 (1931) |
| Multiplicity | 5.7 (5.9) |
| Completeness %) | 98.52 (96.13) |
| Mean I/sigma (I) | 4.24 (1.12) |
| Wilson B-factor | 21.65 |
| R-merge | 0.2059 (1.281) |
| R-meas | 0.2266 (1.406) |
| R-pim | 0.09307 (0.5691) |
| CC1/2 | 0.986 (0.571) |
| CC* | 0.997 (0.853) |
| Reflections used in refinement | 19580 (1913) |
| Reflections used for R-free | 1006 (96) |
| R-work | 0.2040 (0.3152) |
| R-free | 0.2436 (0.3155) |
| CC (work) | 0.944 (0.787) |
| CC (free) | 0.883 (0.716) |
| Number of non-hydrogen atoms | 2356 |
| Macromolecules | 2171 |
| Solvent | 185 |
| Protein residues | 274 |
| RMS (bonds) | 0.008 |
| RMS (angles) | 0.85 |
| Ramachandran favored (%) | 98.12 |
| Ramachandran allowed (%) | 1.88 |

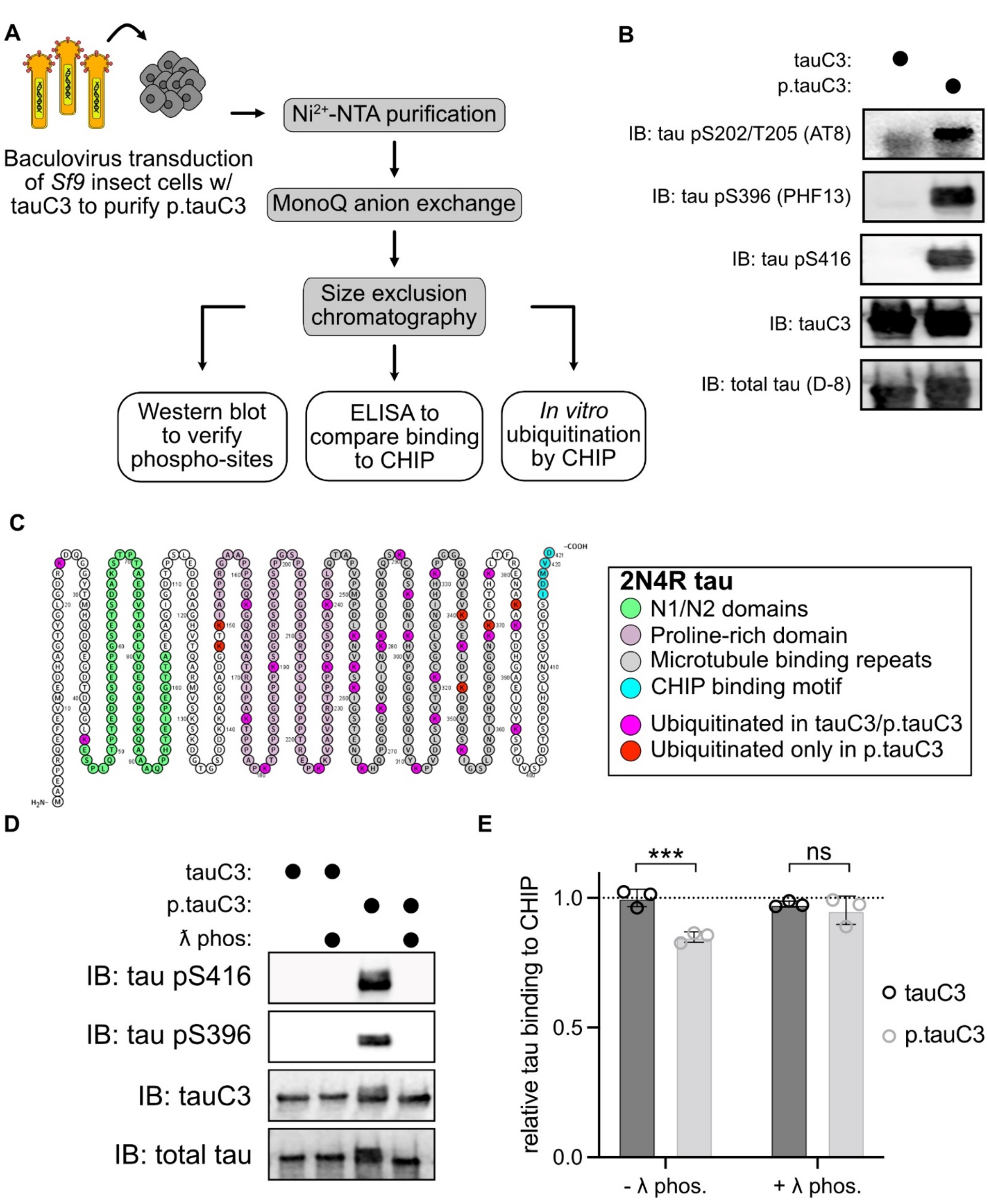


**Supplemental Fig 1. Expression of tauC3 in insect cells yields protein that is phosphorylated at multiple sites, including pathogenic residues.** **(A)** Cartoon depicting workflow for purification of p.tauC3 from *Sf9* insect cells and subsequent analyses. **(B)** Western blot confirmation of phospho-epitopes on p.tauC3. tauC3 and total tau antibodies were used as loading controls. **(C)** Cartoon depicting identified sites of ubiquitination on tauC3 or p.tauC3 following *in vitro* ubiquitination by CHIP. Shared sites for tauC3 and p.tauC3 are depicted in purple, while sites unique to p.tauC3 are shown in red. **(D)** Western blot confirming dephosphorylation of p.tauC3 by λ phosphatase. **(E)** TauC3 or p.tauC3 binding to immobilized CHIP following dephosphorylation by λ phosphatase measured by ELISA. Assay was performed in triplicate and normalized to tauC3 absorbance at 450 nM without λ phosphatase treatment. Statistical significance was determined by two-way ANOVA with Bonferroni's post-hoc analysis (ns = not significant, ***$p<0.001$, n = 3).

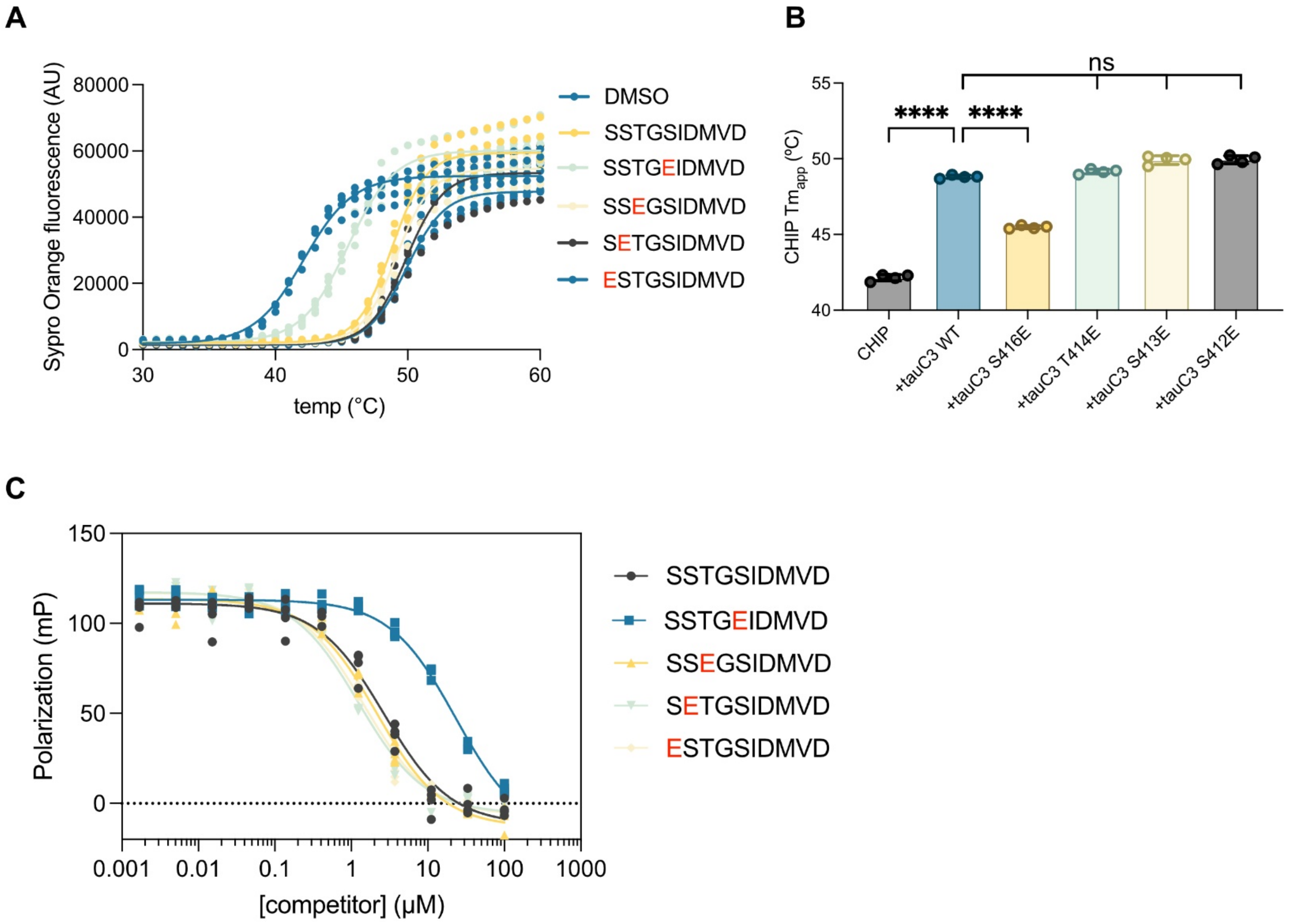


**Supplemental Fig. 2 Pseudo-phosphorylation at serine 416 is sufficient to inhibit CHIP binding. (A)** DSF melt curves for CHIP in the presence of various 10-mer tauC3 phosphomimetic peptides. Assay was performed in quadruplicate and melt curves were fit with a Boltzmann sigmoid. **(B)** Apparent melting temperatures ($Tm_{app}$) of CHIP WT in the absence or presence of 10-mer tau peptides as derived from (A). Statistical significance was determined by one-way ANOVA with Tukey's post-hoc analysis (ns = not significant, ****$p<0.0001$, n = 4). **(C)** Competition FP experiment showing displacement of fluorescent tracer from the CHIP TPR domain by various 10-mer tau peptides. Samples were performed in quadruplicate.

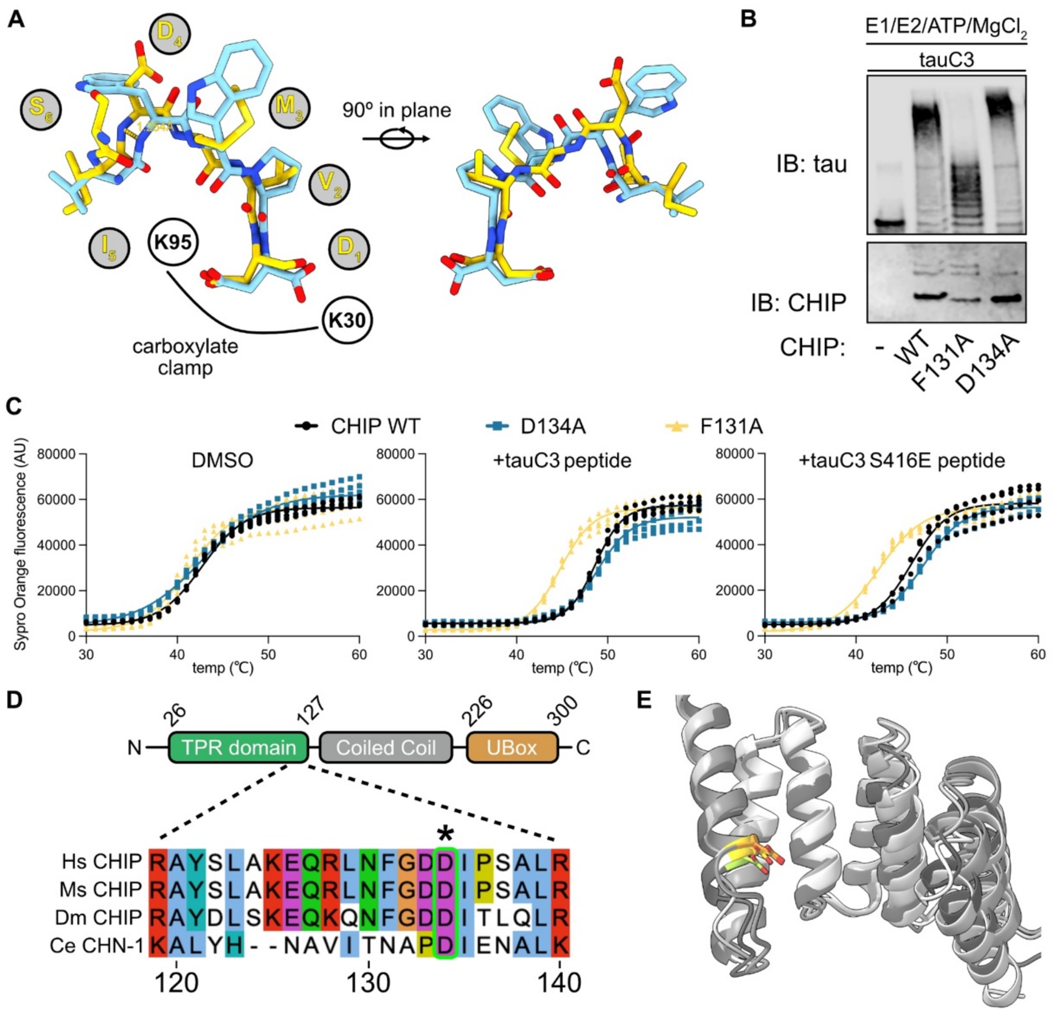


**Supplemental Fig. 3 Mutation of a conserved residue in CHIP (CHIP D134A) partially restores activity on tauC3 pS416. (A)** Structural alignment of tauC3 peptide (yellow) with an optimized CHIP peptide (blue) bound to the CHIP TPR (PDB: 6NSV). Residues 10-7 from the tauC3 peptide are omitted for clarity. TauC3 residues 1-6 are noted in gray circles. Location of the CHIP carboxylate clamp residues are denoted in white open circles. The 1.254 Å shift in the backbone register is shown on the overlay. **(B)** *In vitro* ubiquitination of tauC3 by various CHIP mutants. A single timepoint was collected, quenched in SDS-PAGE loading buffer, and analyzed by western blot. **(C)** DSF melt curves for various CHIP mutants in the absence or presence of various 10-mer tauC3 peptides. Assay was performed in quadruplicate and melt curves were fit with a Boltzmann sigmoid. **(D)** Cartoon depicting the domain architecture of the CHIP monomer, with region of the TPR domain bearing D134 highlighted below. Conservation of D134 across evolution is highlighted in green with asterisk. (Hm = *H. sapiens*; Ms =*M. musculus*; Dm =*D. melanogaster*; Ce = *C. elegans*). **(E)** Structural overlay of CHIP TPR domain across evolution. AlphaFoldv2.0 structures from human, mouse, fruit fly, and nematode CHIP are shown in various shades of gray, with D134 highlighted in various shades of yellow.

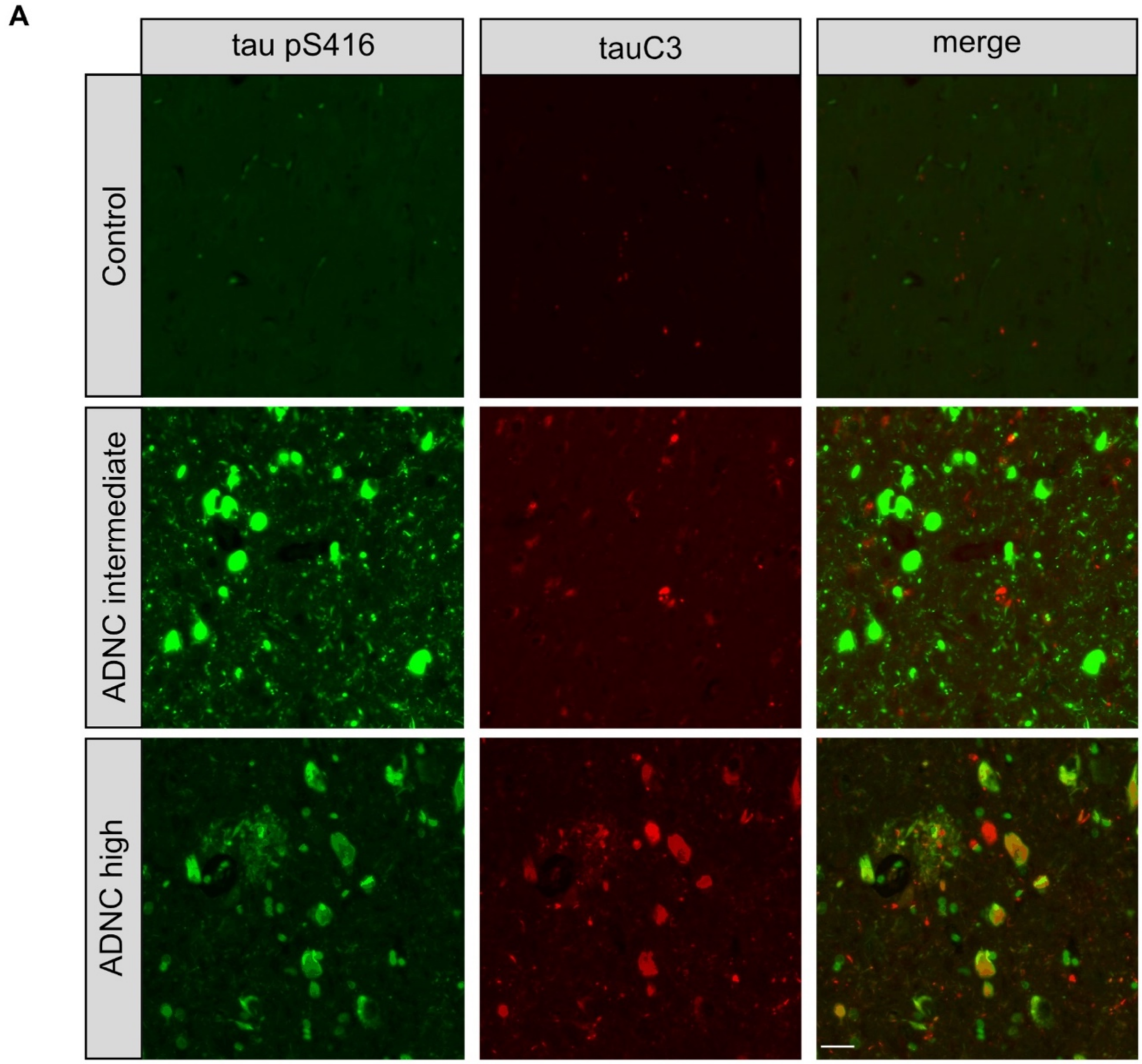


**Supplemental Fig. 4 Additional examples of pS416 accumulation in AD and co-localization with tauC3.** Representative immunofluorescence micrographs from the subiculum of human patient samples across increasing ADNC score. Tau pS416 is shown in green, while tauC3 staining is shown in red. Scale bar = 50 μM.
